# Cooperativity of weak actomyosin interaction

**DOI:** 10.1101/2024.05.28.596264

**Authors:** Aarushi Naskar, Alexis Johnson, Yuri E. Nesmelov

**Affiliations:** Department of Physics and Optical Science, UNC Charlotte, Charlotte NC 28223; Department of Biological Sciences, UNC Charlotte, Charlotte NC 28223

**Author notes:** These authors contributed equally to this work (Aarushi Naskar, Alexis Johnson).

**Keywords:** myosin, actin, muscle, cooperativity, weak interaction

## Abstract

We report the discovery of a new regulatory mechanism of the actomyosin system in muscle. We show that the weak binding of the myosin-nucleotide complex with unregulated F-actin is a cooperative process. Hundreds of myosin heads must work together for efficient force production in muscle, but the precise mechanism by which they coordinate remains elusive. It is known that myosin initially binds actin weakly, then transitions into a strongly bound state to produce force. Using the contiguous cooperative binding model, we interpreted our experimental results in terms of a cooperativity parameter defined as an increased probability for a myosin head to bind to the actin filament next to the already bound head. Considering the geometric organization of a sarcomere, we propose the formation of cross-bridge clusters composed of up to six myosin heads bound consecutively to actin. The cooperativity of weak actomyosin interaction may explain several challenging questions in muscle physiology, such as the role of myosin isoforms in mixed-isoform hybrid muscles, or the yield of supramaximal rate of force production in decorated skinned muscle fibers.

**Significance Statement:** Force in striated muscle results from myosin interacting with actin. Initially, myosin attaches weakly to the thin filament, transitioning to a strongly bound state, generating force. Our experiments show high cooperativity in myosin’s weak interaction with unregulated actin filament. This cooperative behavior may facilitate the formation of cross-bridge clusters and the cooperative steps of myosin heads between clusters. Consequently, the thin- and thick-filament regulation could govern the spacing between cross-bridge clusters and influence the probability of a myosin head stepping along the thin filament during force development in muscle.

## Introduction

A sarcomere, the force-producing unit of muscle, consists of parallel thin and thick filaments. The filaments slide relative to each other when muscle contracts and myosin heads, tethered to the thick filament, bind actin of thin filament, and produce ATP-fueled conformational change. The current hypothesis of the force production in muscle considers myosin heads as individual force producers; coordination of myosin heads in the force production is not assumed.

Muscle force production is regulated by troponin, the part of the troponin-tropomyosin regulatory complex of thin filament, controlling availability of myosin binding sites. Calcium, released in response to external signaling, changes troponin conformation, opening seven-to-eleven myosin binding sites on thin filament (1, 2). The relationship between calcium concentration in muscle and produced force is highly cooperative. Currently it is assumed that cooperativity is governed by the three-step mediation of actomyosin interaction by troponin-tropomyosin regulatory complex of thin filament (3) and that the cooperativity is lost at full calcium activation of thin filament, when all binding sites are open (2). The majority of studies examined myosin interaction with unregulated F-actin or thin filaments used myosin in the post-power stroke state and assumed non-cooperative weak actomyosin interaction. Myosin head has three structural states in muscle: the active state when the head is bound to thin filament (post power stroke structural state M), the disordered relaxed state (DRX) when the head is ready to bind thin filament (nucleotide bound, pre power stroke M** state), and the super relaxed (storage) state when the head is parked on thick filament sequestered from actin. In actomyosin cycle, ADP release follows ATP-induced actomyosin dissociation and the head transitions into the DRX state. The lifetime of myosin head in the DRX state can reach tens of seconds (4).

Because physiological concentration of ATP in muscle is in the millimolar range, the majority of myosin heads are in the DRX state, available for force production. Myosin transitions from the DRX state by binding thin filament initially weakly and then strongly. Transient kinetics of weak actomyosin interaction when myosin is in the M** structural state was examined in several studies and the reported data are controversial. Marston and Taylor (5) observed approximately exponential transient of actomyosin interaction in the double mixing experiment when myosin heads were first premixed with ATP and then mixed with unregulated F-actin. White and Taylor (6) and Stein et al (7, 8) reported sigmoidal transient of weak actomyosin interaction in their experiments. The sigmoidal transient is indicative of the cooperativity of the weak interaction of myosin and unregulated F-actin. Following previous studies, we examined interaction of myosin heads in the M** structural state with unregulated F-actin in detail and used the model of contiguous binding of ligand to one-dimensional polymer (9) to interpret our transient kinetics data. We conclude that myosin head prepared in the pre-power stroke structural state M** binds unregulated F-actin cooperatively and the mechanism of cooperative force development in muscle should include the cooperativity of the weak binding of myosin head and actin of thin filament.

## Materials and Methods

### Ethical approval

Myosin S1 and actin were produced from rabbit skeletal tissue. All experimental protocols were approved by the Institutional Animal Care and Use Committee of UNC Charlotte and all preparations were performed in accordance with relevant guidelines and regulations.

### Reagents

Phalloidin, ATP, and ADP were from Sigma-Aldrich (Milwaukee, WI). All other chemicals were from ThermoFisher Scientific (Waltham, MA) and VWR (Radnor, PA).

### Preparation of proteins

Myosin and actin acetone powder were prepared from rabbit leg and back muscles (1). Full length myosin was digested with chymotrypsin (30 mg/L) for 10 minutes at T=20°C, the reaction was terminated with 2mM phenylmethylsulfonyl fluoride. Undigested myosin was removed by centrifugation and myosin S1 was dialyzed in the experimental buffer.

Actin was prepared from acetone powder by three consecutive repolymerization cycles using 50 mM KCl and 2 mM MgCl_2_. F-actin was stabilized with phalloidin with the molar ratio 1:1. Concentration of G-actin and myosin S1 was determined spectrophotometrically assuming the extinction coefficient ε_290nm_ = 0.63 (mg/ml)^-1^cm^-1^ for actin and ε_280nm_ = 0.83 (mg/ml)^-1^cm^-1^ for myosin (1, 2). The experimental buffer contained 50 mM Tris-HCl pH 7.5, 50 mM KCl, 3mM MgCl_2_, 100 μM CaCl_2_, 1mM DTT. All reported concentrations are final concentrations. All experiments were performed with the freshly prepared myosin S1. F-actin stabilized with phalloidin was stored at +4°C and used no later than three weeks after preparation.

### Acquisition of transients

We used Bio-Logic SFM-300 stopped-flow transient fluorimeter (Bio-Logic Science Instruments SAS, Claix, France), equipped with an FC-15 cuvette and thermostated syringe and cuvette compartments. To characterize kinetics of myosin-ATP interaction, single mixing scheme was used. Myosin solution was rapidly mixed with MgATP solution and transient change of myosin intrinsic fluorescence was detected by the photomultiplier tube. The wavelength of excitation light was 302 nm and emitted light passed 320 nm cut-off filter. To characterize rigor myosin-actin binding, myosin solution was rapidly mixed with F-actin solution and transient change of light scattering at 400 nm was detected by the photomultiplier tube. In the double-mixing experiments, equal volumes of myosin and MgATP solutions of equal concentrations were rapidly mixed, and the resulting mixture was transferred into the delay line with a volume of 90 μL. A triple volume of the mixed solution was transferred through the delay line to wash it. Following a controlled delay, the solution from the delay line was rapidly mixed with the actin solution, and the flow was stopped to acquire transient changes in light scattering, reflecting weak actomyosin interaction. A flow rate of 8 ml/s was utilized in all transient experiments. Multiple transients were acquired and averaged to improve the signal-to-noise ratio.

The instrument’s dead time was independently calibrated with MgCl_2_ and hydroxyquinoline (3), resulting in a value of 2.6 ms. A total of 8000 points were acquired in each experiment.

### Analysis of fluorescence transients

To characterize myosin-ATP interaction and rigor actomyosin interaction obtained transients were fitted by the single exponential function S(t) = S_o_+A·exp(-k_obs_·(t-t_0_)). S(t) is the observed signal at the time t, A is the signal amplitude, t_0_ is the time before the flow stops, and k_obs_ is the observed rate constant. In the case of myosin-ATP interaction, the dependence of the observed rates on the ATP concentration was fitted by a hyperbola, v = V_max_·[ATP]/(K_app_ +[ATP]), allowing the determination of the maximum rate, V_max_ (the horizontal asymptote) (Figure S4). To determine the bimolecular rate K_1T_k_+2T_ the dependence of the observed rates on the ATP concentration was fitted by a straight line at small concentrations of ATP. In the case of the rigor actomyosin interaction the dependence of observed rate constants on myosin S1 concentration was fitted with a straight line to determine the averaged rate (Figure S5). These experiments were conducted routinely to characterize myosin and actin preparations.

Transients of weak actomyosin interaction were analyzed in terms of the contiguous binding model. Obtained transients were fitted to the numerical solution of Eq. 21 using Monte-Carlo least squares fitting approach. Three parameters were extracted from the fits, the forward and reverse rate constants of weak actomyosin interaction k_ON_ and k_OFF_ and the cooperativity parameter ω. In our analysis we assumed cooperative actomyosin binding and non-cooperative actomyosin dissociation. Rate constants were obtained in experiments with myosin S1 and actin from multiple independent preparations. Obtained rate constants were averaged and mean values and standard deviations are reported. Data fits with standard functions such as an exponent, hyperbola, and line were performed with Origin Pro (OriginLab Corp, Northampton MA).

### Statistical analysis

The data reported as mean value ± SD. For mean data, the number of observations (N) corresponds to the number of independent protein preparations. In protein preparations we used muscle tissue from three rabbits. Obtained transients were fitted to the contiguous binding model using Monte-Carlo routine and the least squares method. The coefficient of determination (R^2^) was used to characterize the goodness of fit for all fitted curves, the fit was considered good and was accepted when R^2^ > 0.99. We used two-sample t-test analysis to examine if the kinetic constants and cooperativity parameter obtained from fits of transients acquired at different temperatures are statistically different. The p-value was calculated to determine if the null hypothesis (equal kinetic constants and cooperativity parameter) or the alternative hypothesis (not equal kinetic constants and cooperativity parameter) is true. We assumed that the p-value less than the significance level of 0.05 indicates that the null hypothesis can be rejected.

Variances of different datasets were compared using two-sample test for variance with the null hypothesis that the datasets have equal variances vs. the alternative hypothesis that the datasets do not have equal variances. Statistical analysis was performed using the statistics package integrated into Origin (OriginLab Corp, Northampton MA) software, assuming that the kinetic constants are independent and normally distributed.

### The model

The model for the contiguous binding of ligands to a one-dimensional polymer was initially developed for steady-state interaction by McGee and von Hippel (5) and later adapted for transient kinetics assays by Villaluenga et al. (6). The model analyzes the probability of occupancy of two consecutive binding sites on a polymer (Figure 1). Defining the pair of two consecutive binding sites on the polymer as *(ff)* when both sites are free on the left and right, and *(bf)* and *(fb)* when a ligand is bound to the left site of the pair and the right site is free, and vice versa, and *(bb)* when both the left and right sites of the pair have bound ligands, one can write the following expressions,

**Figure 1.**
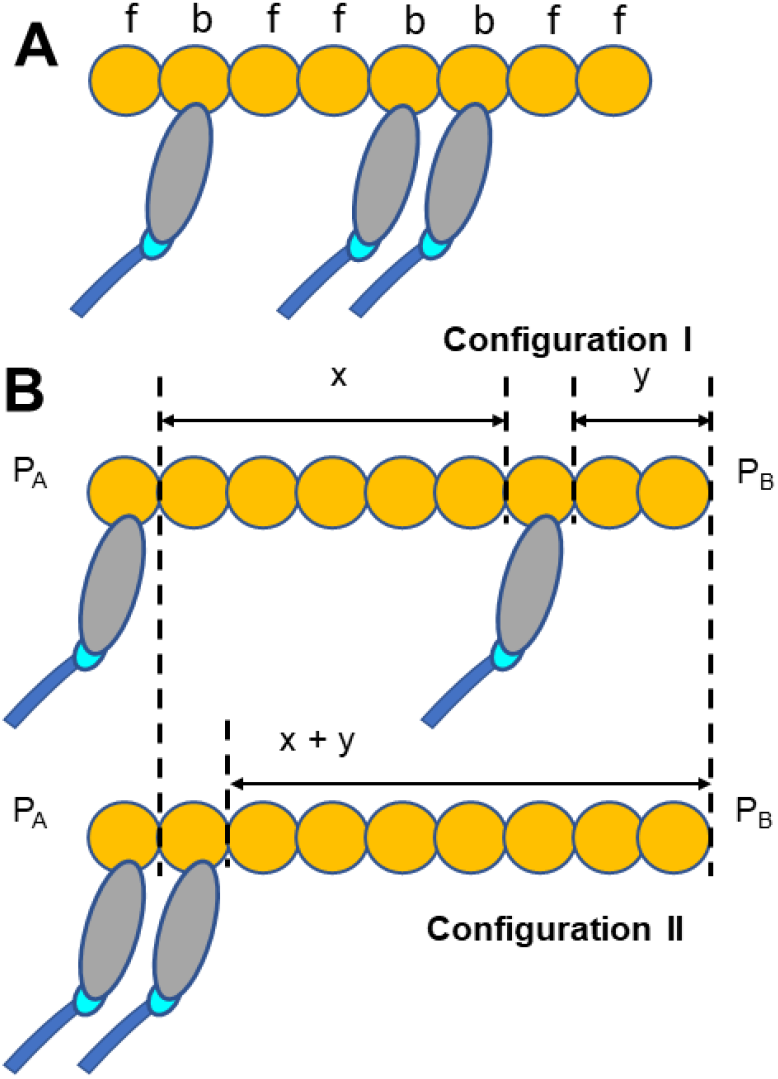
Schematic illustration of polymer and ligand consecutive binding pairs (A), and the definition of the cooperativity parameter as the ratio of configuration II and configuration I (B). “f” and “b” indicate free and occupied binding sites of the polymer, respectively. In (A) the following pairs of consecutive binding sites are depicted from left to right, (fb), (bf), (ff), (fb), (bb), (bf), (ff). In (B) “P_A_” and “P_B_” denote the same probabilities of configurations of bound ligands on the left and right of the considered configurations. Conditional probabilities of configurations I and II reflect non-cooperative and cooperative ligand binding, respectively.

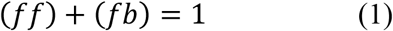

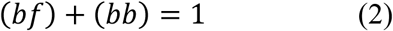

Equation (1) illustrates that a pair with a free left site can either have a free or bound site on the right, and the sum of these conditional probabilities for the pair is equal to one. The same logic applies to Equation (2). The occupancy of the left site of the pair can be described by the following equation,

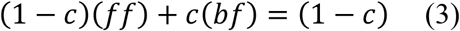

where the coefficient *c* represents the coverage. The occupancy of the binding sites in a pair is considered random, and in this equation, the term *(1−c)* denotes the probability of having the left site of the pair free, while the term *c* represents the probability of having the left site of the pair occupied. This equation is equivalent to

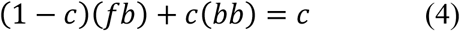

which describes the probability of finding a free or occupied left site in the pair with the occupied right site. The cooperativity parameter *ω* is defined as the ratio of probabilities of two configurations of bound ligands, cooperative and non-cooperative, as illustrated in Figure 1.

Assuming that in both cases the probabilities of flanking configurations of bound ligands on the left (*P*_*A*_) and right (*P*_*B*_) are the same, the probability of non-cooperative configuration (configuration I in Figure 1) is

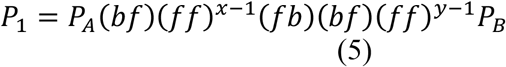

and the probability of the cooperative case of ligand binding configuration (configuration II in Figure 1) is

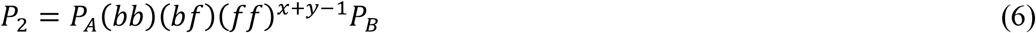

Then the cooperativity parameter defined through the conditional probabilities is

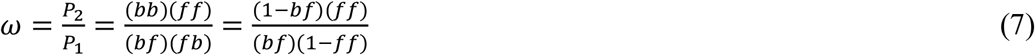

Assuming a large number of myosin binding sites in an actin polymer, *N*, with *n* myosin heads bound, resulting in *n+1* gaps between bound heads, the average number of free binding sites per gap, or the expected value *s*, is given by

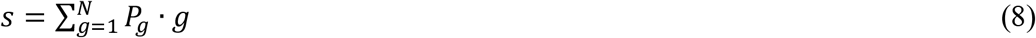

where *P*_*g*_ is the probability that any particular gap has *g* consecutive free binding sites. One can see that *P*_1_ = (*bf*)(*fb*), *P*_2_ = (*bf*)(*ff*)(*fb*), …, *P*_*g*_ = (*bf*)(*ff*)^*g*−1^(*fb*). For the non-cooperative case, conditional probabilities *(bf)* and *(ff)* are equal because the probability to bind a free site on the right in a pair does not depend on the occupancy of the left site. Then, for a long polymer, the number of free binding sites per gap is

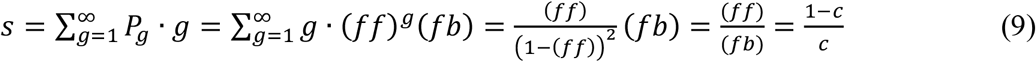

where 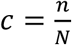.

The differential equation describing the kinetics of ligand binding to a polymer is

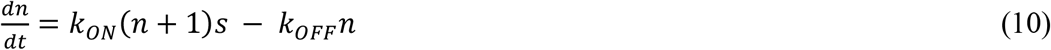

and in the non-cooperative case, it simplifies to

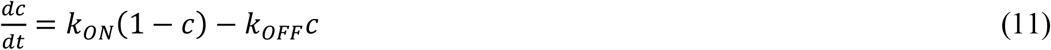

as expected, assuming n ≅ n+1 for a long polymer.

For the cooperative case, conditional probabilities *(bf)* and *(ff)* are not equal because the probability to bind a free site on the right in a pair depends on the occupancy of the left site. One can use Eq. S7 to find the relation between the conditional probabilities *(bf)* and *(ff)*,

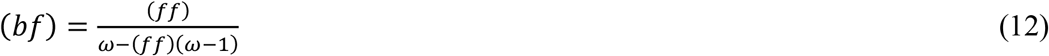

Starting from Eq. (3) and Eq. (7), simple but tedious rearrangement allows to express conditional probability *(ff)* in terms of coverage *c* and cooperative parameter *ω*,

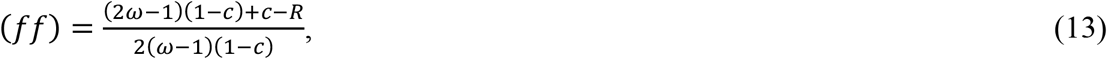

where

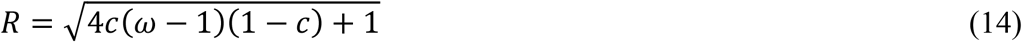

Therefore,

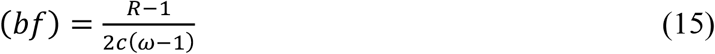

There are three types of free binding sites in a gap: doubly contiguous, singly contiguous, and individual binding sites. The expected value for the number of double contiguous binding sites per gap is

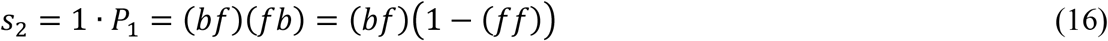

The expected value for two singly contiguous binding sites per gap is

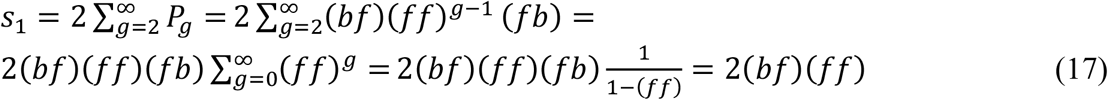

The expected value for the individual binding sites in a gap is

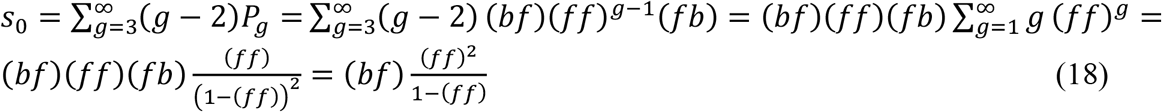

The total expected value can be expressed as a sum

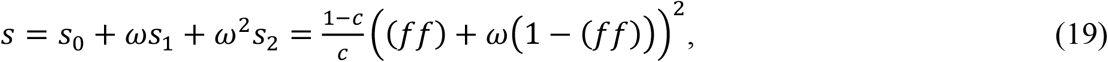

And therefore, using Eq. (10),

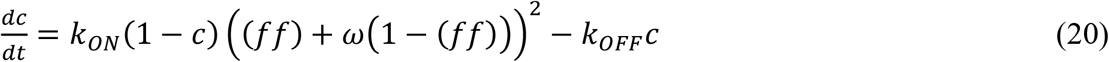

Combining Eq. (20) and Eq. (13) one can obtain following kinetic equation, accounting for individual, single, and double contiguous binding of myosin heads to actin filament in our experiment

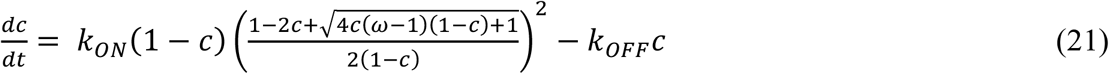

## Results

To study the kinetics of weak actomyosin interaction, we conducted a double mixing transient kinetics experiment. Myosin S1 was prepared in the pre-power stroke state M** by rapidly mixing equimolar amounts of myosin S1 and ATP, with the time of M** state formation determined in a separate experiment (Figure S1). We selected a 2-second incubation time for our double mixing experiment, changing the delay time in the 0.5 s – 10 s range did not affect the results. Following a controlled two-second delay, M** myosin S1 was rapidly mixed with unregulated F-actin. After the flow stops, the transient change in light scattering of the final actomyosin mix reflects the binding of myosin S1 and actin (Figure 2). Similar transients were observed when monitoring the fluorescence of pyrene-labeled actin.

**Figure 2.**
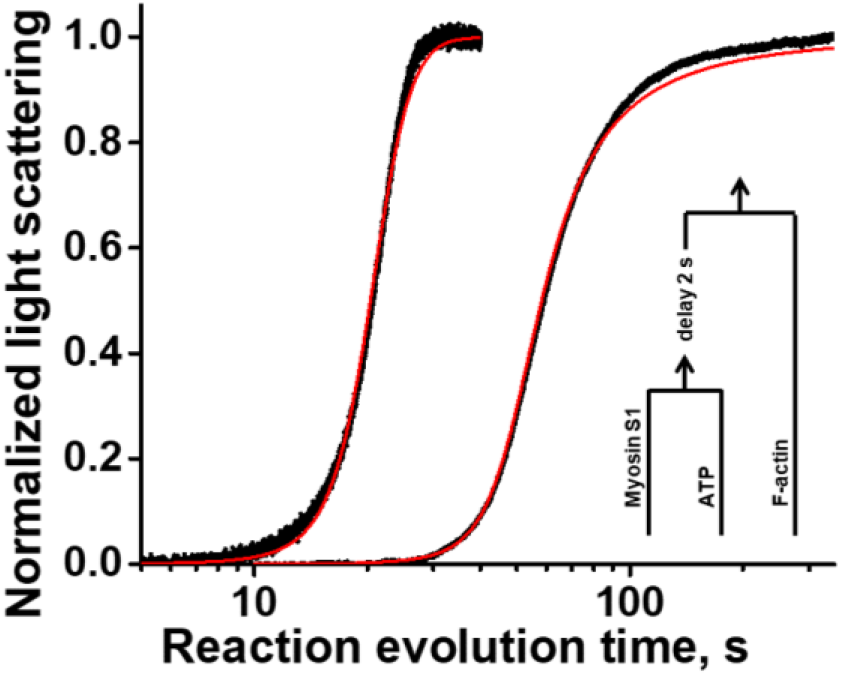
Representative traces of the weak actomyosin interaction. Myosin S1, premixed with equimolar ATP and incubated for two seconds in a delay line, was rapidly mixed with F-actin and transient of light scattering was observed. Left transient, [myosin]=[ATP]=5μM, [actin]=0.5μM. Right transient, [myosin]=[ATP]=[actin]=1μM. All concentrations are final. Both transients fitted simultaneously with the same set of reaction rates k_ON_, k_OFF_, and the cooperativity parameter ω. Black dots, obtained transients, red trace – simultaneous fit. T=20oC

All observed transients exhibit a lag followed by a rise in light scattering (Figure 2). One interpretation of such transients could be that during the lag phase, myosin hydrolyzes ATP and subsequently binds to actin. An alternative interpretation of these transients involves the cooperative binding of myosin S1 to actin filaments, wherein the rate of myosin S1 binding next to already bound myosin head exceeds the rate of binding to bare actin filaments. To distinguish between these interpretations, we conducted a double mixing experiment where actin filaments decorated with rigor myosin S1 were used in the second mixing stage. We anticipated that if there is no cooperativity, there would be no change in the transient’s position. Conversely, if cooperativity exists, the transient should shift to the left, as myosin already bound to actin filament would promote the weak binding of myosin prepared in the M** structural state. Figure 3 shows that decorating actin filaments with rigor myosin S1 shifts the transient to the left, with the magnitude of the shift depending on the concentration of decorating myosin S1. This experiment leads us to conclude that weak actomyosin interaction is cooperative.

**Figure 3.**
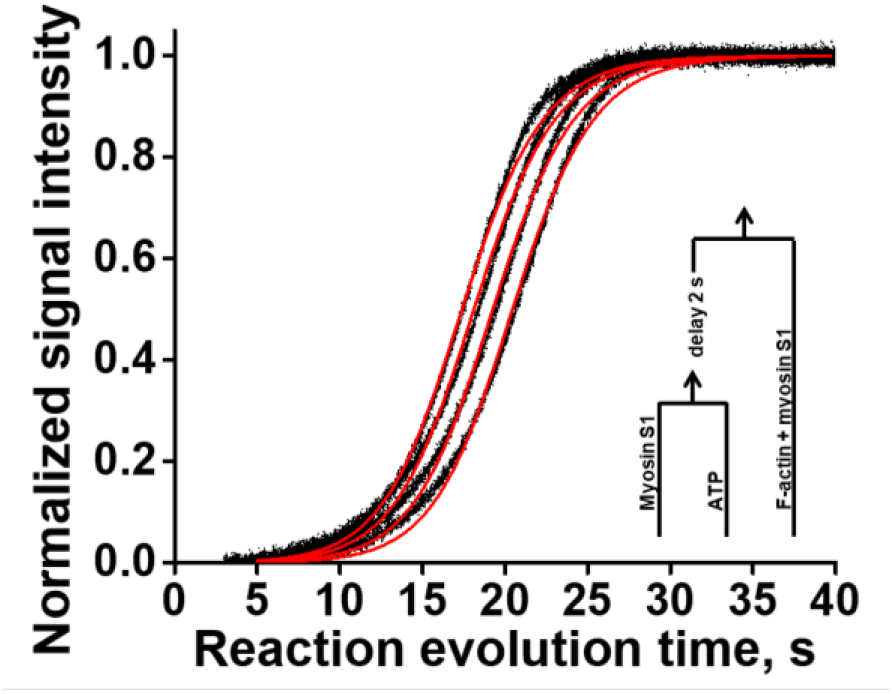
An example of the weak interaction of myosin S1 with decorated F-actin. Myosin S1, premixed with equimolar ATP to form the M** structural state, follows into a delay line and after a two second delay rapidly mixed with F-actin, decorated with myosin S1 of different concentrations. Final concentrations are 5μM myosin M**, 0.5μM actin, and concentrations of decorating myosin S1 are 0μM, 0.1μM, 0.25μM, and 0.5μM, right to left accordingly. Assuming equilibrium association constant of rigor myosin–actin interaction as 9.1 μM^-1^, final concentrations of decorating myosin are 0μM, 0.08μM, 0.19μM, and 0.31μM. T=20°C.

The shape of a transient is determined by the evolution of the reaction mixture and is described by a solution of the corresponding differential equation. In the case of non-cooperative binding, such as the rigor binding of myosin S1 and unregulated F-actin (Figure S2), the simple reaction scheme 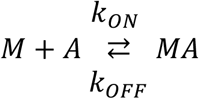 is described by the differential equation [*MA*]^′^ = −*k*_*ON*_[*M*][*A*] + *k*_*OFF*_[*MA*], where *M* and *A* stand for myosin and actin, and *k*_*ON*_ and *k*_*OFF*_ are the reaction rate constants. To account for the cooperativity effect, we fitted observed transients to the model of contiguous binding of ligand to a one-dimensional polymer (Eq. 21). In the model, the rate constant of the weak actomyosin binding, *k*_*ON*_, is modulated by the ligand coverage of a polymer. The transients shown in Figures 2 and 3 were fitted with the model of contiguous binding using the same set of kinetic parameters, reaction rates and cooperativity parameter, but with a varied concentration of myosin S1 in the M** state (Figure 2), and varied fraction of rigor myosin S1 of the decorated actin filament (Figure 3). In Figure 3 the shift of the transient can be explained as an increased probability of weak actomyosin interaction when the “seed” myosin heads are already in place on the actin filament and serve as promoters of the binding. Since the equilibrium constant of rigor actomyosin binding is well known (10, 11), one can estimate the fraction of bound myosin heads of the decorated actin filament using the following equation,

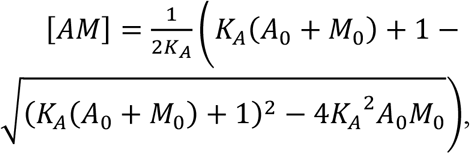

which is the meaningful solution of the quadratic equation

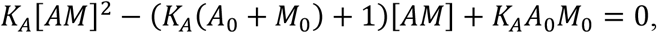

obtained from the expression for the equilibrium association constant *K*_*A*_ = [*AM*]/[*A*][*M*] and actin and myosin concentrations, *A*_0_ = [*A*] + [*AM*] and *M*_0_ = [*M*] + [*AM*], where A_0_ and M_0_ are the total actin and myosin concentrations used in the experiment, and [*A*], [*M*], and [*AM*] are concentrations of actin, myosin, and actomyosin in the reaction mixture. We used 0.1 μM, 0.25 μM, and 0.5 μM myosin S1 to decorate actin filaments (0.5 μM, all final concentrations), which correspond to 0.08 μM, 0.18 μM, and 0.31 μM assuming *K*_*A*_ = 9.1 μM ^-1^ (10). Our fits show a thousand times smaller number of “seed” heads on the decorated actin filament, 0.04±0.02 nM, 0.14±0.06 nM and 0.28±0.01 nM accordingly (Figure S3). It is possible that for cooperative binding, the “seed” myosin heads should be in a certain stereospecific orientation on the actin filament, and not all rigor cross-bridges on the decorated actin filament are in that state to promote cooperative binding. Bershitsky et al. suggested that in muscle fiber for successful force production myosin heads should be in a specific bound conformation (12). Our experiments confirm the importance of the right orientation of a crossbridge on the actin filament. Previously, it was proposed that bound myosin changes the conformation of actin in the filament to promote the binding of other myosin heads (13). If this is the case, we expect to see a different reaction rate constant *k*_*ON*_ in our experiment, since the cooperativity parameter reflects only the probability of contiguous binding. Our results show that the reaction rate constant *k*_*ON*_ does not change significantly upon actin filament decoration, and therefore, bound myosin does not change the conformation of actin in the filament, promoting binding of other heads. The result, similar to actin filament decoration was observed when we varied the concentration of ATP in the preparation of myosin S1 in the M** state. When the ATP concentration is lower than the concentration of myosin, the transient shifts left, since there is a mixture of myosin heads in the M and M** states, and rigor myosin quickly binds the actin filament, forming “seeds” for the cooperative weak binding. When ATP is in excess, the transient shifts right; the shift can be explained as multiple ATP turnovers before final binding to the actin filament. When during the preparation of myosin heads in the M** state, myosin S1 is premixed with the mixture of ADP and ATP, the transient shifts right due to the competition of ADP and ATP binding to myosin, the same effect as when ATP is in excess upon the preparation of myosin in the M** state. An increase in temperature causes the transient to shift towards the origin and alters its shape (see Figure 4, Table 1). Our analysis demonstrates that reaction rates are influenced by temperature, while the cooperativity parameter remains unaffected, consistent with expectations. However, we note that experiments conducted at T=37°C may yield less reliable results. When heat sample to 37°C and reducing the temperature back to 20°C, we occasionally observed a rightward shift in the transient’s position compared to its position before the temperature increase. We attribute this shift to partial degradation of myosin S1 at 37°C, leading to a decrease in active myosin heads available to interact with the actin filament. Consequently, we excluded data obtained at T=37°C when such a shift occurred upon temperature reduction. Statistical analysis reveals that the variance of the rate coefficient *k*_*ON*_ was similar for T=12°C and T=20°C, and their mean values differed significantly, with a p-value of 0.004. The variance of the coefficient *k*_*ON*_ for T=37°C differed from that of T=20°C and T=12°C. The mean value of *k*_*ON*_ at T=37°C was statistically similar to that of transients acquired at T=12°C and T=20°C, with p-values of 0.25 and 0.3, respectively. The variances and mean values for the coefficient *k*_*OFF*_ and the cooperativity parameter were similar across all tested temperatures. Our study aimed to test the hypothesis of cooperativity in weak actomyosin interactions, and we conducted experiments at various temperatures primarily to confirm the observation of cooperativity across temperature ranges. The investigation into the temperature dependency of weak actomyosin interactions will be conducted separately.

**Table 1.**
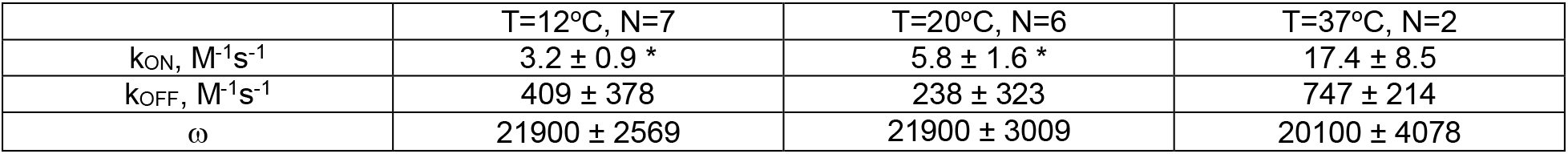
Kinetic rate constants and the cooperativity parameter of weak actomyosin interaction, determined at various temperatures. N denotes the number of biological repeats. For experiments conducted at 37°C, we ensured the reproducibility of transients obtained at 20°C before and after reaching 37°C. If transients at the lower temperature were not reproducible, those data were excluded from the analysis. An asterisk denotes statistically different results.

| | $T=12^{\circ}\text{C}$, N=7 | $T=20^{\circ}\text{C}$, N=6 | $T=37^{\circ}\text{C}$, N=2 |
| --- | --- | --- | --- |
| $k_{ON}$, $\text{M}^{-1}\text{s}^{-1}$ | $3.2 \pm 0.9$ * | $5.8 \pm 1.6$ * | $17.4 \pm 8.5$ |
| $k_{OFF}$, $\text{M}^{-1}\text{s}^{-1}$ | $409 \pm 378$ | $238 \pm 323$ | $747 \pm 214$ |
| $\omega$ | $21900 \pm 2569$ | $21900 \pm 3009$ | $20100 \pm 4078$ |

**Figure 4.**
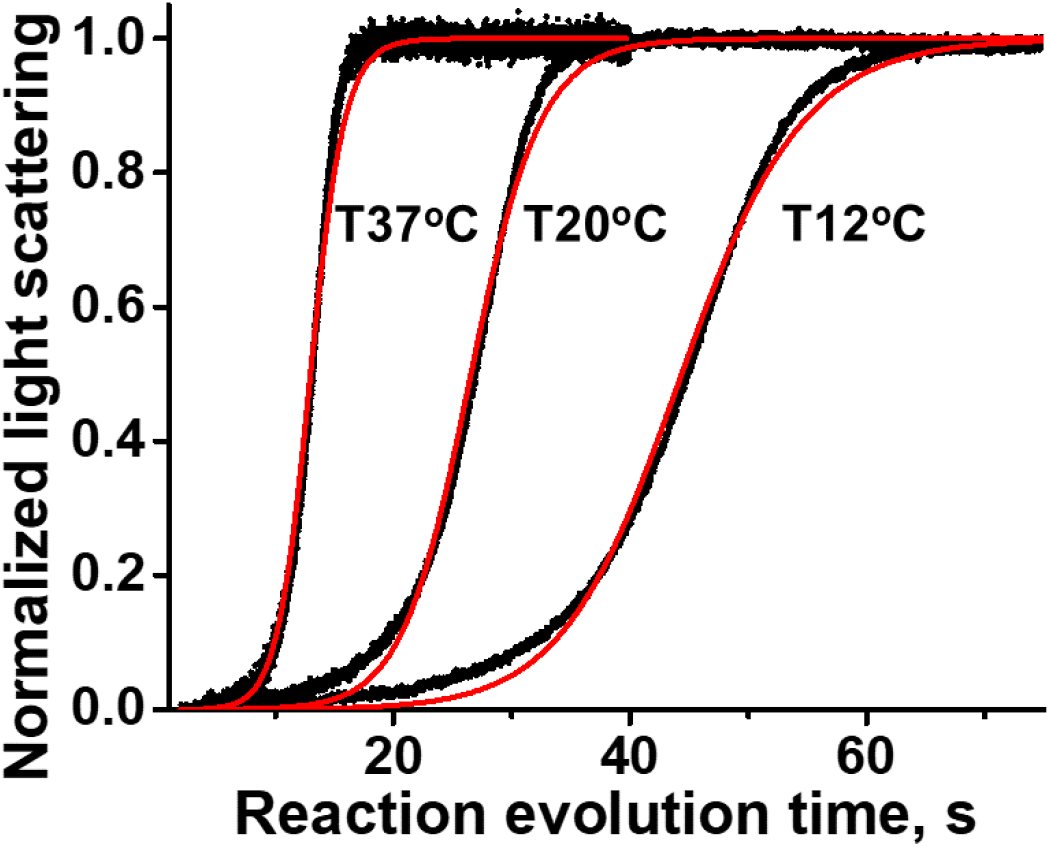
Representative transients of weak actomyosin interaction acquired at various temperatures. The concentrations were held constant, [myosin] = [ATP] = 5 μM and [actin] = 0.5 μM. The corresponding reaction rates k_ON_, k_OFF_, and the cooperativity parameter ω, are presented in Table 1. Black dots, obtained transients, red trace, fit.

## Discussion

The three-state model of the thin filament (3) explains the cooperativity of rigor myosin binding to the thin filament. We argue that there is a difference in actomyosin interaction when myosin is in the M (rigor) or M** (pre-power stroke) state. Our experiments show that the myosin head in the M** state binds unregulated actin filament cooperatively. Rapid mixing of myosin in the M** state with decorated F-actin results in faster kinetics of the weak actomyosin binding, the same as the decoration of skinned muscle fiber with NEM myosin results in the increased rate of force development (14). We propose that the calcium-dependent thin filament regulation of myosin binding and cooperativity of weak actomyosin interaction are complementary. The cooperativity of weak actomyosin interaction may result in the formation of cooperative clusters of cross-bridges bound to the thin filament and the cooperative myosin steps between the clusters upon muscle contraction.

Actomyosin interaction in solution differs from interactions in muscle fiber; in solution, the myosin head is free to bind any available binding site, whereas in the fiber, there are constraints due to the spatial distribution of myosin heads, thin filament binding sites, and constraints from myosin head tethers. Therefore, the size of cooperative clusters of myosin heads on the thin filament cannot be determined from the model using just expected values of the probability of binding. According to the model, when the cooperativity parameter is equal to one, the probability of the availability of singly contiguous binding sites is 50%, and the probabilities to have available double contiguous and individual sites both are 25%. An increase in the cooperativity parameter redistributes these probabilities, and the probability of having available individual sites quickly increases to 90% and beyond, reflecting the formation of large cooperative clusters of myosin heads and significant gaps between them. Obviously, this cannot be the case for muscle fiber. To estimate the size of a cooperative cluster of cross-bridges in a fiber, one can assume that for efficient force production, the myosin head in the M** state always binds cooperatively and steps over a gap between cooperative clusters. For skeletal muscle, the step size is 10 - 28 nm (15), which agrees with the proposed 22 nm long S2 tether of the myosin head (16). Using available structures of myosin heads bound to F-actin (5h53.pdb, 5jlh.pdb), one can estimate the length of the myosin binding site on the actin filament in the axial direction as 5.4 nm. Therefore, the step size of 10 – 28 nm corresponds to a step over one to five actin monomers, meaning that the size of the gap between cooperative clusters is two – six actin monomers. To estimate the length of the cooperative cluster, we need to know the number of myosin heads and actin monomers in the overlapped region of the thick and thin filaments in a half sarcomere. In the sarcomere of skeletal muscle, the thin and thick filaments form a hexagonal structure; one thick filament is surrounded by six thin filaments, and three thick filaments neighbor one thin filament. The thick filament is a bipolar, 1600 nm long filament, containing 294 dimers of myosin heads (17). There is a 300 nm long bare zone in the middle of the thick filament that does not contain myosin heads (18). The length of the myosin heads-containing part of the half-thick filament is 650 nm in a half sarcomere, and the number of myosin heads in the half-thick filament is 294. A thin filament is approximately 1000 nm long, but since only 650 nm of the thick filament contains myosin heads, the 650 nm long segment of the thin filament plays a role in muscle contraction due to the filaments’ overlap. Since each thick filament provides myosin heads for six thin filaments, and there are three thick filaments surrounding one thin filament, the average number of myosin heads per one thin filament is 147. There are 245 actin monomers in the 650 nm long segment of the thin filament (19), which agrees well with our estimate of the linear dimension of the thin filament binding site. Bershitsky et al (12) reported that in the fully activated muscle up to 75% of myosin heads interact with actin, which reduces the number of active myosin heads per thin filament from 147 to 110. This gives the ratio of active myosin heads to actin monomers in the overlap region of the contracting muscle approximately 1:2, and the numbers of occupied and free sites of the thin filament are approximately equal. Therefore, one can expect the cluster size of two to six thin filament-bound myosin heads, separated by two to six free binding sites on a thin filament. We hypothesize that the myosin head steps from the end of one cluster to the beginning of another one in the inchworm manner. We cannot rule out the hand-over-hand myosin steps, but considering the distance between central axes of thick filaments as 35 nm (20, 21), one can calculate the distance between central axes of thick and thin filaments as 23.3 nm. Estimating the distance from the axis of the thin filament to the C-terminus of the relay helix of the catalytic domain of the bound myosin head (the place of connection of the catalytic and converter domains of the myosin head) as 8.3 nm (5h53.pdb), and the radius of the thick filament backbone as 12.5 nm (22), there is not much room left for the hand-over-hand stepping of the myosin head in muscle fiber. If the number of active myosin heads decreases (for example, due to the transition into the super-relaxed (SRX) state (23)), the length of a gap between clusters increases, and the myosin cycle becomes less efficient, since a cycling head must bind the thin filament non-cooperatively. From the perspective of the cooperativity effect, the transition to the SRX state has a regulatory effect, since the size of a gap between cooperative clusters of bound myosin heads depends on the number of available active heads and the coverage of thin filament. It is interesting to estimate how determined rates of weak actomyosin interaction are compared with the rate of steady-state actin-activated myosin ATPase assay and the rate of force redevelopment in skinned skeletal muscle fiber, since both depend on the rate of weak to strong transition of actomyosin interaction. At contiguous cooperative binding the rate of binding is modulated by the coverage (Eq. 21), and therefore not a constant value. We determined the rate of myosin S1 binding to naked actin at temperature 20°C as 5.8 ± 1.6 M^−1^s^−1^ (Table 1). The term, modulating the reaction rate constant *k*_*ON*_ (Eq. 21) has maximum value of 20490 for the nearly complete coverage of actin filament for the determined cooperativity parameter (Table 1). The product of the rate and the modulating term gives the reaction constant 0.12 µM^−1^s^−1^ for cooperative weak actomyosin interaction at T=20°C. At saturating actin concentration of 100 µM, the rate of actin-activated myosin ATPase activity then should be 12 s^−1^, which agrees well with the literature data for rabbit skeletal myosin S1 and actin (24-27). The rate of force redevelopment in skinned rabbit skeletal muscle fiber after 10% slack is 15 s^−1^ (14). Assuming sarcomere length 2.2 µm, the force redevelopment after the 10% slack corresponds to half-sarcomere shortening by 110 nm. As discussed above, if the length of the myosin step is 10 – 28 nm, four to eleven steps of the myosin head on the thin filament are required for force redevelopment. To estimate the rate of one step, we need to know the concentration of myosin heads in the sarcomere. The concentration of myosin in skeletal muscle was determined as 105 µM (28). This corresponds relatively well to a simple estimate, using the number of myosin heads in the half-thick filament, the distance between thick and thin filaments, and the half-sarcomere length. Indeed, assuming 294 myosin heads in a cylinder with a radius equal to the thick-thin filament distance (23.3 nm) and the length of a half-sarcomere (1.1 µm), the concentration of myosin heads in the sarcomere is 260 µM. But considering constraints for the myosin head (the length of the overlap region and unavailability of space occupied by the thick filament backbone), the estimate of the concentration of myosin heads in the sarcomere increases to 600 µM. Using the same rate of cooperative weak actomyosin interaction, 0.12 µM^−1^s^−1^, and the estimated concentration of myosin heads, one can obtain the rate for one step as 72 s^−1^. Since four to eleven steps are required for force redevelopment, the rate should be in the range of 6.5 s^−1^ – 18 s^−1^, which corresponds well to the measured rate of force redevelopment in skinned rabbit skeletal muscle fiber (14). Cooperativity of weak actomyosin interaction can elucidate the role of fast myosin isoform in hybrid muscles as a fast thin filament binder, producing “seeds” for cooperative clusters formation. The cooperative work of myosin isoforms in muscle may increase the time of the strong actomyosin bond in a cycle, which increases muscle power output. The rate of weak actomyosin binding is maximal when binding is cooperative, therefore, modulation of cooperative myosin stepping in sarcomere by changing the availability of binding sites on thin filament or availability of myosin heads to form cooperative cross-bridges has a regulatory effect.

## Author Contributions

YN, designed research, AN and AJ performed research, AJ and YN analyzed data, AN, AJ, YN wrote the paper.

## Acknowledgments

This research was funded by the National Institutes of Health, grant number HL159585.

## References

1. R. Desai, M. A. Geeves, N. M. Kad, Using fluorescent myosin to directly visualize cooperative activation of thin filaments. J Biol Chem 290, 1915–1925 (2015).

2. K. M. Trybus, E. W. Taylor, Kinetic studies of the cooperative binding of subfragment 1 to regulated actin. Proc Natl Acad Sci U S A 77, 7209–7213 (1980).

3. D. F. McKillop, M. A. Geeves, Regulation of the interaction between actin and myosin subfragment 1: evidence for three states of the thin filament. Biophys J 65, 693–701 (1993).

4. C. R. Bagshaw, D. R. Trentham, The reversibility of adenosine triphosphate cleavage by myosin. Biochem J 133, 323–328 (1973).

5. S. B. Marston, E. W. Taylor, Comparison of the myosin and actomyosin ATPase mechanisms of the four types of vertebrate muscles. J Mol Biol 139, 573–600 (1980).

6. H. D. White, E. W. Taylor, Energetics and mechanism of actomyosin adenosine triphosphatase. Biochemistry 15, 5818–5826 (1976).

7. L. A. Stein, P. B. Chock, E. Eisenberg, The rate-limiting step in the actomyosin adenosinetriphosphatase cycle. Biochemistry 23, 1555–1563 (1984).

8. L. A. Stein, R. P. Schwarz, Jr., P. B. Chock, E. Eisenberg, Mechanism of actomyosin adenosine triphosphatase. Evidence that adenosine 5’-triphosphate hydrolysis can occur without dissociation of the actomyosin complex. Biochemistry 18, 3895–3909 (1979).

9. J. D. McGhee, P. H. von Hippel, Theoretical aspects of DNA-protein interactions: co-operative and non-co-operative binding of large ligands to a one-dimensional homogeneous lattice. J Mol Biol 86, 469–489 (1974).

10. A. H. Criddle, M. A. Geeves, T. Jeffries, The use of actin labelled with N-(1-pyrenyl)iodoacetamide to study the interaction of actin with myosin subfragments and troponin/tropomyosin. Biochem J 232, 343–349 (1985).

11. L. E. Greene, E. Eisenberg, Dissociation of the actin.subfragment 1 complex by adenyl-5’-yl imidodiphosphate, ADP, and PPi. J Biol Chem 255, 543–548 (1980).

12. S. Y. Bershitsky et al., Muscle force is generated by myosin heads stereospecifically attached to actin. Nature 388, 186–190 (1997).

13. A. Orlova, E. H. Egelman, Cooperative rigor binding of myosin to actin is a function of F-actin structure. J Mol Biol 265, 469–474 (1997).

14. D. P. Fitzsimons, R. L. Moss, Cooperativity in the regulation of force and the kinetics of force development in heart and skeletal muscles: cross-bridge activation of force. Adv Exp Med Biol 592, 177–189 (2007).

15. T. Q. Uyeda, S. J. Kron, J. A. Spudich, Myosin step size. Estimation from slow sliding movement of actin over low densities of heavy meromyosin. J Mol Biol 214, 699–710 (1990).

16. R. K. Brizendine et al., A mixed-kinetic model describes unloaded velocities of smooth, skeletal, and cardiac muscle myosin filaments in vitro. Sci Adv 3, eaao2267 (2017).

17. S. J. Atkinson, M. Stewart, Molecular basis of myosin assembly: coiled-coil interactions and the role of charge periodicities. J Cell Sci Suppl 14, 7–10 (1991).

18. H. A. Al-Khayat, R. W. Kensler, E. P. Morris, J. M. Squire, Three-dimensional structure of the M-region (bare zone) of vertebrate striated muscle myosin filaments by single-particle analysis. J Mol Biol 403, 763–776 (2010).

19. T. D. Pollard, Measurement of rate constants for actin filament elongation in solution. Anal Biochem 134, 406–412 (1983).

20. T. Irving et al., Thick-filament strain and interfilament spacing in passive muscle: effect of titin-based passive tension. Biophys J 100, 1499–1508 (2011).

21. B. C. Tanner, T. L. Daniel, M. Regnier, Sarcomere lattice geometry influences cooperative myosin binding in muscle. PLoS Comput Biol 3, e115 (2007).

22. T. F. Robinson, L. Cohen-Gould, Myofilament diameters: an ultrastructural re-evaluation. Adv Exp Med Biol 170, 47–61 (1984).

23. J. W. McNamara, A. Li, C. G. Dos Remedios, R. Cooke, The role of super-relaxed myosin in skeletal and cardiac muscle. Biophys Rev 7, 5–14 (2015).

24. J. M. Chalovich, L. A. Stein, L. E. Greene, E. Eisenberg, Interaction of isozymes of myosin subfragment 1 with actin: effect of ionic strength and nucleotide. Biochemistry 23, 4885–4889 (1984).

25. S. S. Margossian, S. Lowey, Preparation of myosin and its subfragments from rabbit skeletal muscle. Methods Enzymol 85 Pt B, 55–71 (1982).

26. S. S. Rosenfeld, E. W. Taylor, The ATPase mechanism of skeletal and smooth muscle acto-subfragment 1. J Biol Chem 259, 11908–11919 (1984).

27. R. F. Siemankowski, M. O. Wiseman, H. D. White, ADP dissociation from actomyosin subfragment 1 is sufficiently slow to limit the unloaded shortening velocity in vertebrate muscle. Proc Natl Acad Sci U S A 82, 658–662 (1985).

28. U. Murakami, K. Uchida, Contents of myofibrillar proteins in cardiac, skeletal, and smooth muscles. J Biochem 98, 187–197 (1985).

