## Supplemental Information for "Cooperativity of weak actomyosin interaction"

**Supporting figures, mentioned in the main text**

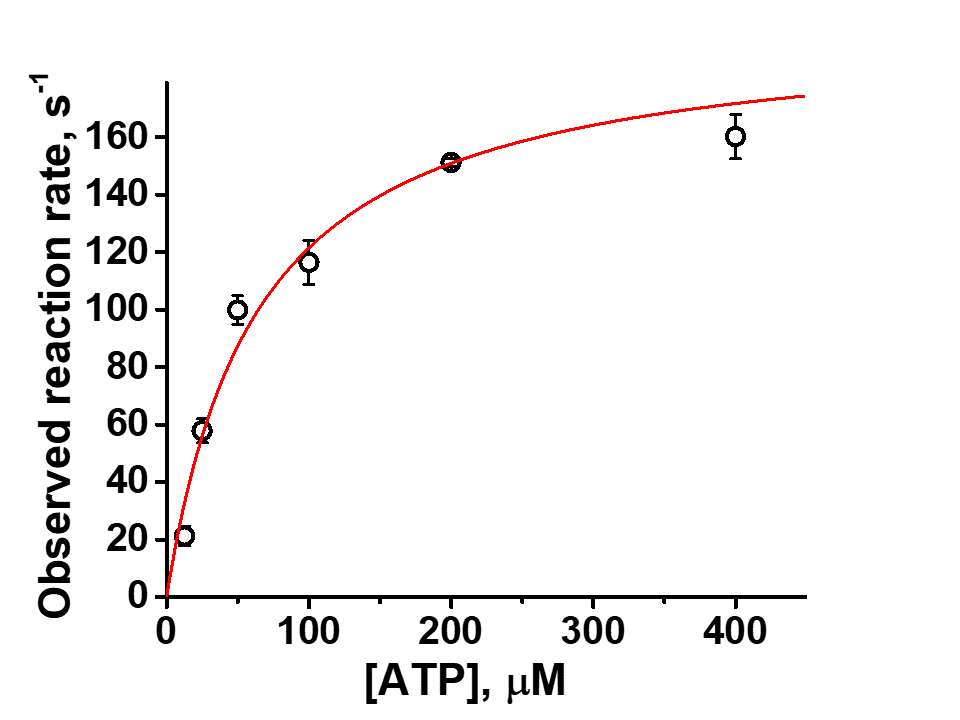

**Figure S 4.** The relationship between the observed rate constant of association of 2 μM myosin S1 with ATP. N=3, T=20°C. The fit with a hyperbola gives v_max_ = 199 s^-1^ and K_D_ = 64 μM. K_1_k_+2_ = 2.1 μM^-1^ s^-1^.

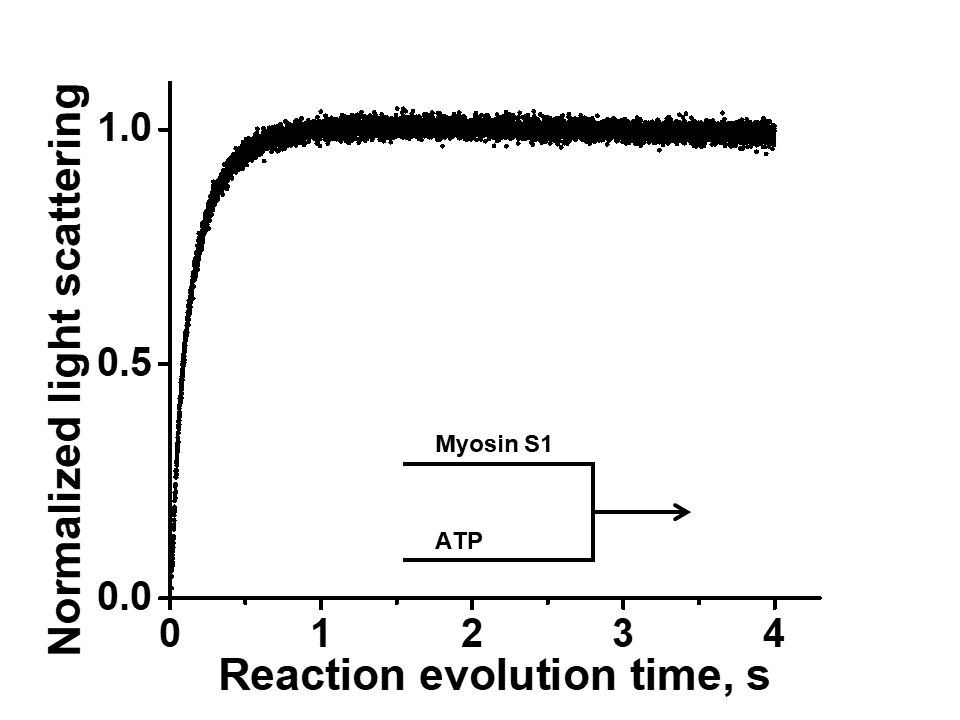

**Figure S 1.** Rabbit muscle myosin S1 is rapidly mixed with ATP, and myosin intrinsic fluorescence reflects conformation change from the post-powerstroke (M) to the pre-powerstroke (M**) state. Myosin quickly adopts M** conformation and slow release of products of ATP hydrolysis eventually brings myosin back to the post-powerstroke (M) structural state. [S1]=5μM, [ATP]=5μM. The max amplitude of the transient corresponds to the max amplitude of the transient when ATP is In excess, indicating that all myosin is in M** structural state after two-second delay. T=20^o^C.

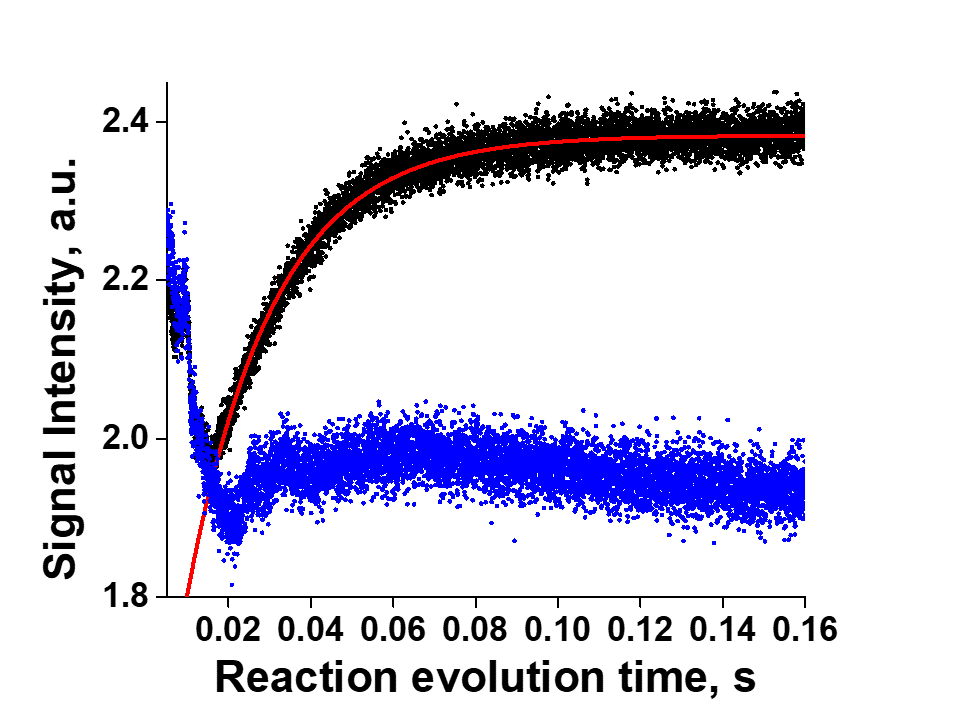

**Figure S 2.** Transients of rigor (black) and weak (blue) actomyosin interaction. Myosin solution is premixed with buffer (rigor binding) or equimolar ATP (weak binding), and after two second delay mixed with actin. The kinetics of the strong rigor binding is described by the exponential function (red trace). Transients of the weak binding usually show some intermediate processes, and then the sigmoidal curve at the later time (Figure 2). Final concentrations, myosin 5 uM, ATP 5 μM, actin 0.5 μM. T=20^o^C.

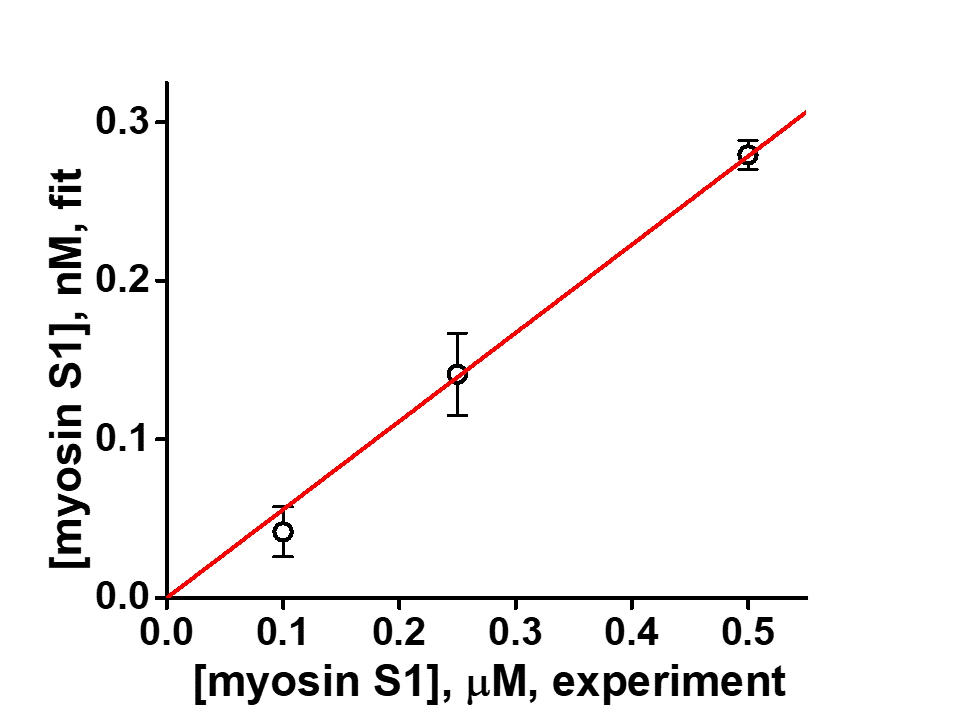

**Figure S 3.** Correlation between the concentration of myosin S1 added to decorate actin in the experiment (x-axis) and the concentration of myosin S1 obtained from fits to the model of cooperative weak actomyosin interaction (y-axis). The slope of the line is 0.56⋅10^-3^, indicating that the concentration of myosin, which acts as a "seed" for cooperative binding, is three orders of magnitude lower than the concentration of myosin added to decorate actin. N=3.

**References**

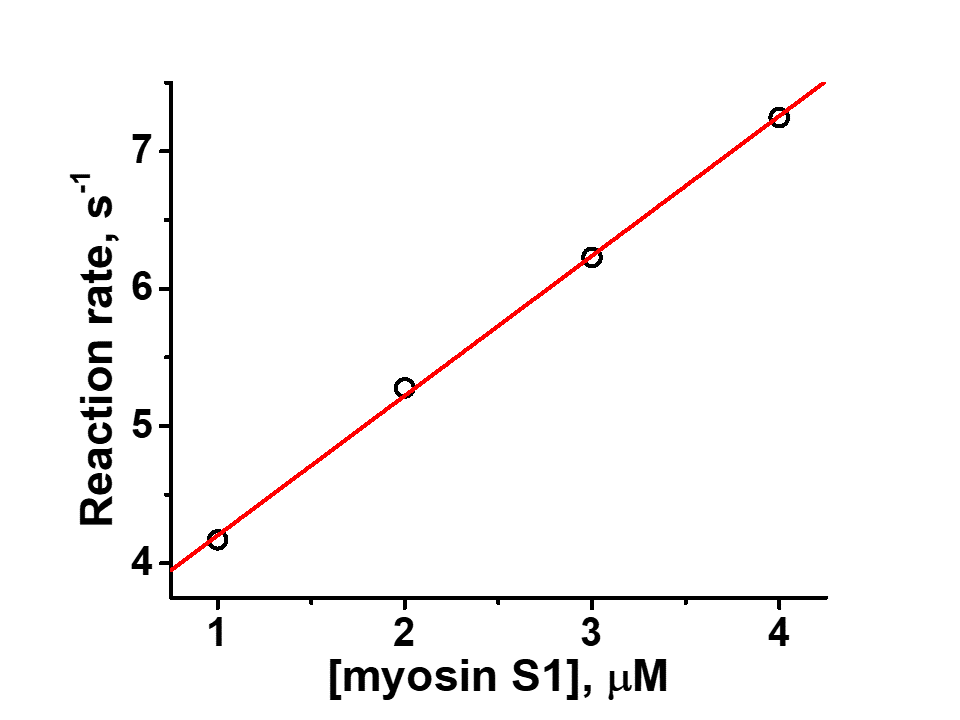

**Figure S 5.** The relationship between the observed rate constant of association of 0.5 μM F-actin with myosin S1 at T=12°C. The slope of the linear fit yields the binding constant k_+_ = 3.19 μM^-1^ s^-1^, in excellent agreement with previous results (1)

1.	S. B. Marston, The rates of formation and dissociation of actin-myosin complexes. Effects of solvent, temperature, nucleotide binding and head-head interactions. *Biochem J* **203**, 453-460 (1982).
